# SIK3 links synaptic strength to sleep pressure through RhoA signaling

**DOI:** 10.64898/2026.09.26.754585

**Authors:** Tomohiro Kitazono, Motoki Juichi, Kei Nishida, Shinnosuke Nomura, Chika Shimizu, Ami Kato, Yusuke Iino, Natsuki Higashide, Kazuya Miyanishi, Shinya Nakata, Natsumi Ageta-Ishihara, Takeshi Sawada, Shoi Shi, Hiromasa Funato, Masashi Yanagisawa

## Abstract

Sleep is regulated by homeostatic sleep pressure, which accumulates during wakefulness, triggering sleep and dissipates during sleep, leading to wakefulness^1–4^. Although recent evidence suggests that synaptic potentiation encodes sleep pressure, the intracellular mechanisms linking synaptic strength to sleep pressure remain unclear. Here we demonstrate that the sleep-promoting kinase SIK3 links synaptic strength and sleep pressure through RhoA signalling. We found that SIK3 is required for dendritic spine enlargement induced by chemical long-term potentiation (cLTP) in cultured cortical pyramidal neurons and positively regulates spine size in the prefrontal cortex in vivo. cLTP reduced RhoA activity in an SIK3-dependent manner during spine enlargement, whereas activation of RhoA reduced EEG delta density in vivo during non-rapid eye movement sleep (NREMS), an indicator of sleep pressure. SIK3 substrate screening identified several RhoA regulators including GEF-H1, and further biochemical analyses found that phosphorylation by SIK3 activates GEF-H1. However, contrary to our expectation, RhoA facilitation through pharmacological activation of GEF-H1 enlarged spine size in cultured neurons, and expression of constitutively active GEF-H1 increased NREMS delta density in a RhoA-dependent fashion. Together, these results indicate that SIK3 enlarges spine size and enhances NREMS delta density through two RhoA-dependent pathways, driving mutually opposing intermediate phenotypes. Our findings identify two clashing SIK3-RhoA signalling pathways that couple synaptic regulation to sleep homeostasis.

## Introduction

All animals with a nervous system alternate between wakefulness and sleep under homeostatic control^5–7^. However, the mechanisms that implement sleep homeostasis, or the neurophysiological basis of homeostatic “sleep pressure”, remains unresolved^8,9^. We previously identified the serine/threonine kinase SIK3 as a key regulator of sleep^10^. Gain-of-function *Sik3* mutant (*Sleepy*) mice exhibit increased non-rapid eye movement sleep (NREMS) amount and electroencephalogram (EEG) delta density during NREMS, a reliable electrophysiological readout of sleep pressure, whereas reducing SIK3 kinase activity decreases NREMS delta density^11–14^. These findings raise the key question of how SIK3-dependent signalling is translated into cellular-level changes that promote sleep. Synaptic strength has been proposed to encode sleep pressure^15,16^, and our recent study has demonstrated that synaptic potentiation in prefrontal cortex (PFC) excitatory neurons causally increases both NREMS amount and EEG delta density^2^. Moreover, the *Sleepy* mutation causes widespread changes in the phosphorylation and expression of synaptic proteins^11,17^. We therefore hypothesised that SIK3 regulates sleep pressure by controlling synaptic strength.

## Results

### SIK3 facilitates dendritic spine enlargement

Excitatory synaptic strength in PFC pyramidal neurons has been proposed to encode sleep pressure^2^, and SIK3 activity in excitatory neurons positively controls sleep pressure^11^. Furthermore, the size of dendritic spines strongly correlates with excitatory synaptic strength^18–20^. Therefore, we quantified spine size in PFC excitatory neurons in *Sleepy* mice exhibiting high sleep pressure (Figure 1A). We sparsely labelled neurons in PFC^2^ and automatically quantified spine size using DeepD3, a deep-learning framework^21^ (Figure S1 and Methods). As expected, spine size was significantly increased in *Sleepy* mice compared with WT mice, although spine density remained unchanged (Fig.1B and 1C). These results suggest that SIK3 signaling facilitates spine enlargement in excitatory neurons, consequently increasing sleep pressure.

**Figure 1.**
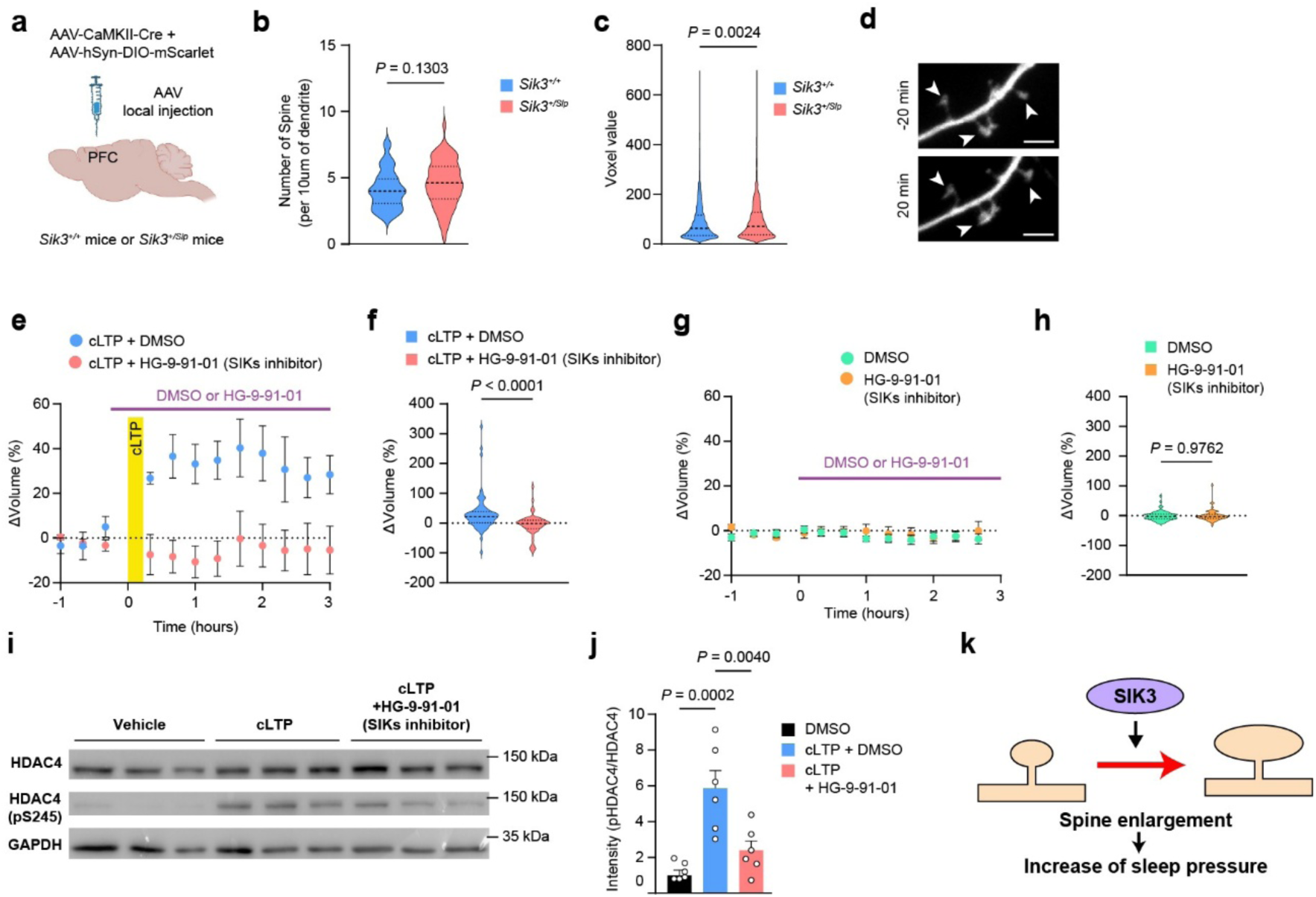
SIK3 regulates dendritic spine size. (A) Scheme of the AAV local injection. (B) The number of spines at ZT12. (*Sik3^+/+^*; n(spines/dendrites/mice) = 298/21/2, *Sik3^Sleepy/+^*; n(spines/dendrites/mice) = 312/20/2, U = 958). (C) The size of spines at ZT12. (*Sik3^+/+^*; n(spines/dendrites/mice) = 1880/54/4, *Sik3^Sleepy/+^*; n(spines/dendrites/mice) = 1908/56/3, U = 1691447). (D) Representative images showing mScarlet before (−20 min) and after (20 min) application of cLTP. Scale bars, 2.5 um. (E) The time course of changes in spine volume under cLTP condition and cLTP with HG-9-91-01 condition. Delta are plotted as a percentage change from the baseline (1 hour before cLTP administration) average (cLTP; n (spine, neuron) =72, 11, cLTP + HG-9-91-01; n (spine, neuron) = 63, 6, U = 1127). (F) The average delta after 20 min after cLTP administration. (G) The time course of changes in spine volume in the presence or absence of HG-9-91-01. Delta are plotted as a percentage change from the baseline (1 hour before drug administration) average (DMSO; n (spine, neuron) = 65, 9, HG-9-91-01; n (spine, neuron) = 66, 9, U = 2138). (H) The average delta after 20 min after drug administration. (I) Representative result of western immunostaining of cultured cortical neurons homogenates under cLTP condition and cLTP with HG-9-91-01 condition. (J) Quantitation of pHDAC4(S245) over HDAC4 band intensity. (K) The proposed role of SIK3 in the regulation of sleep pressure. SIK3 enhances sleep pressure through spine enlargement. Statistical significance was determined by Mann-Whitney U test (B, C, F, H) or one-way ANOVA with Dunnett’s test (J). All error bars correspond to mean ± SEM.

To further examine the role of SIK3 in spine enlargement in cultured cortical neurons, we investigated the effect of SIK3 inhibition during chemical long-term potentiation (cLTP) with a cocktail of bicuculline and glycine in an Mg^2+^-free medium^22^. We added the SIK inhibitor HG-9-91-01^11,17,23^ during cLTP stimulation and quantified spine size. Whereas cLTP stimulation enlarged spines in mock-treated controls (Fig.1d), application of HG-9-91-01 prevented this enlargement (Figure 1D–1F). In contrast, application of HG-9-91-01 without cLTP stimulation did not affect spine size (Figure 1G and 1H). These results suggest that SIK3 activity during cLTP stimulation is required for spine enlargement. Consistent with this, cLTP stimulation enhanced Histone deacetylase 4 (HDAC4) phosphorylation at serine 245 (S245), a canonical SIK3 phosphorylation site^24^, whereas application of HG-9-91-01 suppressed this hyperphosphorylation (Figure 1I and 1J). Taken together, these results support an essential role for SIK3 in cLTP-induced spine enlargement, which may contribute to increased sleep pressure. (Fig.1K).

### SIK3 enlarges spines through RhoA

Although most SIK targets reported to date are transcription regulators^25^, we hypothesized that non-nuclear downstream signaling pathways are involved in synaptic regulation. To explore novel downstream signaling pathways, we performed kinase-oriented substrate screening (KIOSS)^26,27^ in N41 embryonic mouse hypothalamic cells. For SIK3-targeted KIOSS (SIK3-KIOSS) (Figure 2A), global phosphorylation was elevated using the PP1/PP2A phosphatase inhibitor calyculin A, whereas SIK3 activity-sensitive phosphorylation was specifically reduced by co-treatment with HG-9-91-01. The phosphorylation level was measured by LC–MS/MS following 14-3-3 protein– based phosphoprotein enrichment. These pharmacological conditions elevated phosphorylation globally while selectively suppressing the HDAC4 S245 phosphorylation site (Figure S2), thereby validating the SIK-KIOSS screening conditions. SIK-KIOSS identified 584 phosphorylation sites across 204 proteins, of which 32 proteins exhibited a greater than 5-fold reduction in phosphorylation upon co-treatment with HG-9-91-01 (Fig.2B, Table S1 and S2). Pathway analysis of these 32 candidate proteins highlighted Rho GTPase-related pathways, with GEF-H1, ARHGEF7, PLEKHG5, TIAM1, and ARHGAP23 among the candidate proteins (Figure 2B, Table S3). These are either Rho guanine nucleotide exchange factors (GEFs), which activate Rho GTPases, or Rho GTPase-activating proteins (GAPs), which inhibit them, and the majority of them target RhoA, a member of the Rho GTPase family. Based on these findings, we focused on RhoA as a leading candidate factor downstream of SIK3.

**Figure 2.**
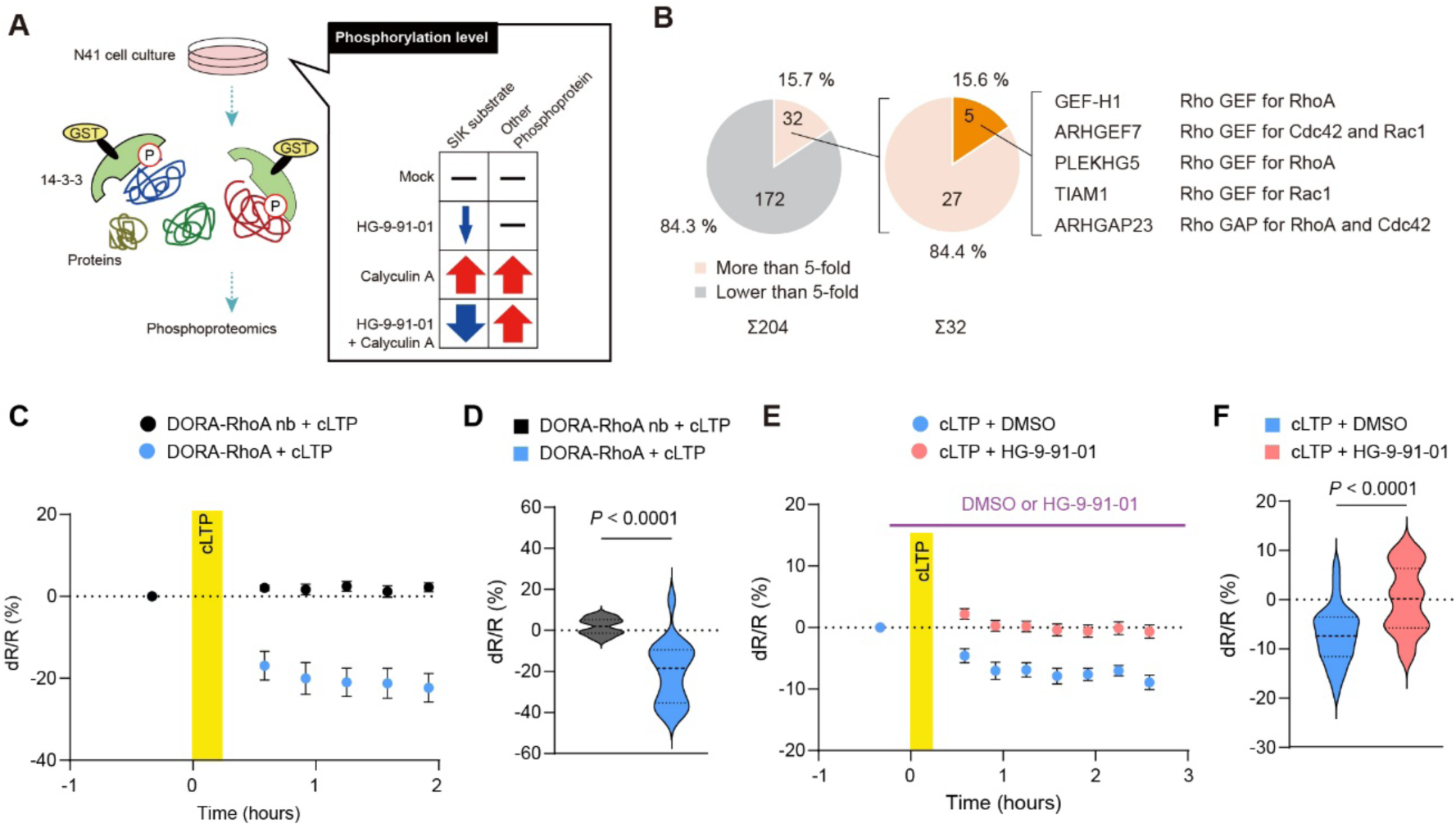
SIK3 enlarges dendritic spine size through RhoA. (A) Scheme and chart for the phosphoproteomic analysis used to identify SIK3 substrates. (B) Pie chart showing the proportion of detected proteins in phosphoproteome analysis (left), pie chart showing the proportion of SIK3 substrate candidates (center), and list of Rho GTPase signaling pathway-related proteins included among the candidates (right). (C) Temporal FRET changes in RhoA activity during cLTP. (D) The average dR/R from 0 to 100 min after cLTP administration (DORA-RhoA nb; n (dendrite, neuron) = 16, 6, DORA-RhoA; n (dendrite, neuron) = 18, 7, U = 149). (E) Temporal FRET changes in RhoA activity during cLTP in the presence or absence of HG-9-91-01. (F) The average dR/R from 0 to 100 min after cLTP administration (DMSO; n (dendrite, neuron) = 30, 9, HG-9-91-01; n (dendrite, neuron) = 55, 16, U = 3860). Statistical significance was determined by Mann-Whitney U test. All error bars correspond to mean ± SEM.

Since RhoA is known to regulate dendritic spine size^28^, we examined whether RhoA functions downstream of SIK3 during spine enlargement in cultured neurons. RhoA activity in spines during cLTP stimulation was visualized with Dimerization Optimized Reporter for Activation (DORA)-RhoA, a RhoA FRET biosensor^29,30^. When cLTP was induced in cultured cortical neurons, the FRET ratio indicating RhoA activity in spines significantly decreased following stimulation, whereas the negative-control mutant sensor showed no change (Figure 2C and 2D). This decrease in RhoA activity was no longer observed in the presence of HG-9-91-01 (Figure 2E and 2F). These results suggest that SIK3 signaling facilitates cLTP-induced spine enlargement by reducing RhoA activity in spines.

### RhoA activation reduces sleep pressure

Since both spine enlargement and synaptic potentiation have been linked to increased sleep pressure^2,15,16^, we next hypothesized that SIK3 and RhoA activities may exert opposing effects on the sleep phenotype. To examine this possibility, we expressed a constitutively active form of RhoA (*Rhoa^Q63L^*) pan-neuronally in adult mice using an AAV vector. As predicted, EEG delta density during NREMS was significantly reduced in RhoA^Q63L^-expressing mice over 24 hours (Figure 3E-3H). Although RhoA^Q63L^ expression did not alter the total amounts of wakefulness, NREMS, or rapid eye movement sleep (REMS), or NREMS episode duration (Figure 3A-3D), it significantly reduced the decreasing time constant (*T*_d_) of Process S, with a resultant reduction in the *T*_d_/*T*_i_ ratio (Figure 3I-3L). These findings indicate that constitutive activation of RhoA selectively reduces the indices of sleep pressure without altering sleep amount.

**Figure 3.**
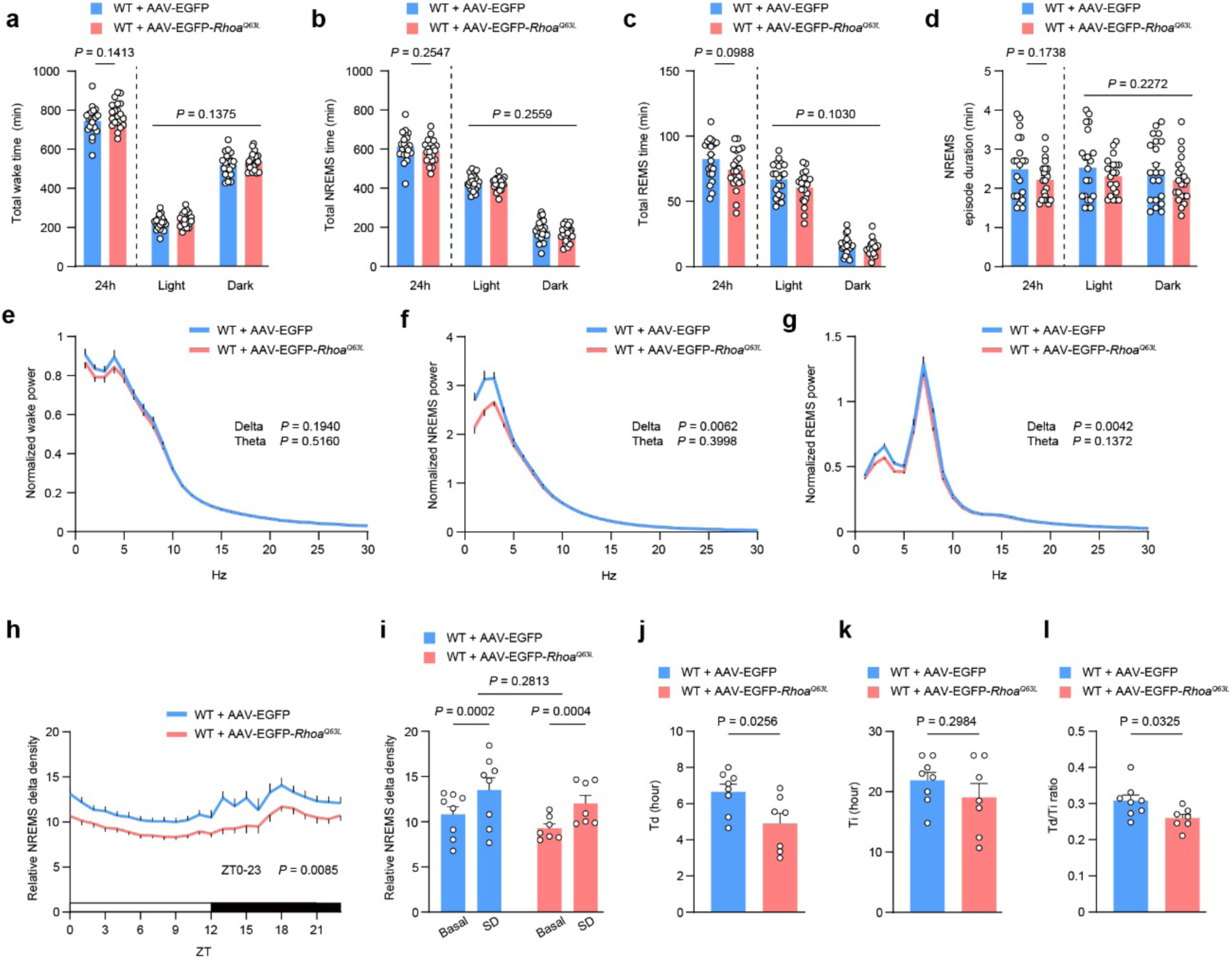
RhoA reduces sleep pressure. (A-D) Total time spent during wakefulness (A), NREMS (B), REMS (C), and NREMS episode duration (D) over 24 hours and during light and dark phases of EGFP expressing or RhoA^Q63L^ expressing mice (EGFP, n = 21; RhoA^Q63L^, n = 21). (E-H) EEG power spectra during wakefulness (E), NREMS (F), REMS (G) over 24 hours and time course of NREMS delta density over 24 hours (H) (EGFP, n = 21; RhoA^Q63L^, n = 21). (I) NREMS delta density during ZT6–ZT8 under the basal condition and after 6-hour sleep deprivation (EGFP, n = 8; RhoA^Q63L^, n = 7). (J-L) *T*_d_ (J), *T*_i_ (K), and the *T*_d_/*T*_i_ ratio (L) derived from Process S fitting for sleep pressure dynamics (EGFP, n = 8; RhoA^Q63L^, n = 7). Statistical significance was determined by two-way ANOVA with Tukey’s test (A-D right, I), unpaired t-test (A-D left, E-G, J-L), or two-way ANOVA with mixed-effects model (H). All error bars correspond to mean ± SEM.

We next examined whether reducing RhoA activity produced the reciprocal phenotype. Brain-wide, pan-neuronal expression of three independent shRNAs for RhoA knockdown reduced RhoA expression in the cortex and hypothalamus by ∼50 % compared with non-targeting shRNAs (Figure S3A). However, this maneuver did not alter the total amounts or spectral profiles of sleep and wakefulness, NREMS delta density after sleep deprivation, or the inferred dynamics of Process S (Figure S3). Similarly, pan-neuronal expression of the dominant-negative mutant RhoA (*Rhoa^T19N^*) resulted in no detectable changes in sleep-wake phenotypes (Figure S4). The absence of sleep phenotypes following RhoA inhibition may reflect incomplete suppression, or functional redundancy and compensation by other members of the Rho GTPase subfamily, such as RhoB and RhoC.

These results demonstrate that constitutive RhoA activation is sufficient to reduce NREMS delta density, producing an effect opposite to the increased NREMS delta density observed in gain-of-function *Sik3* (*Sleepy*) mice^31^, and thereby support a model in which SIK3-dependent reduction of RhoA signaling contributes to the increase in dendritic spine size and sleep pressure indexed by NREMS delta density (Figure 4P).

**Figure 4.**
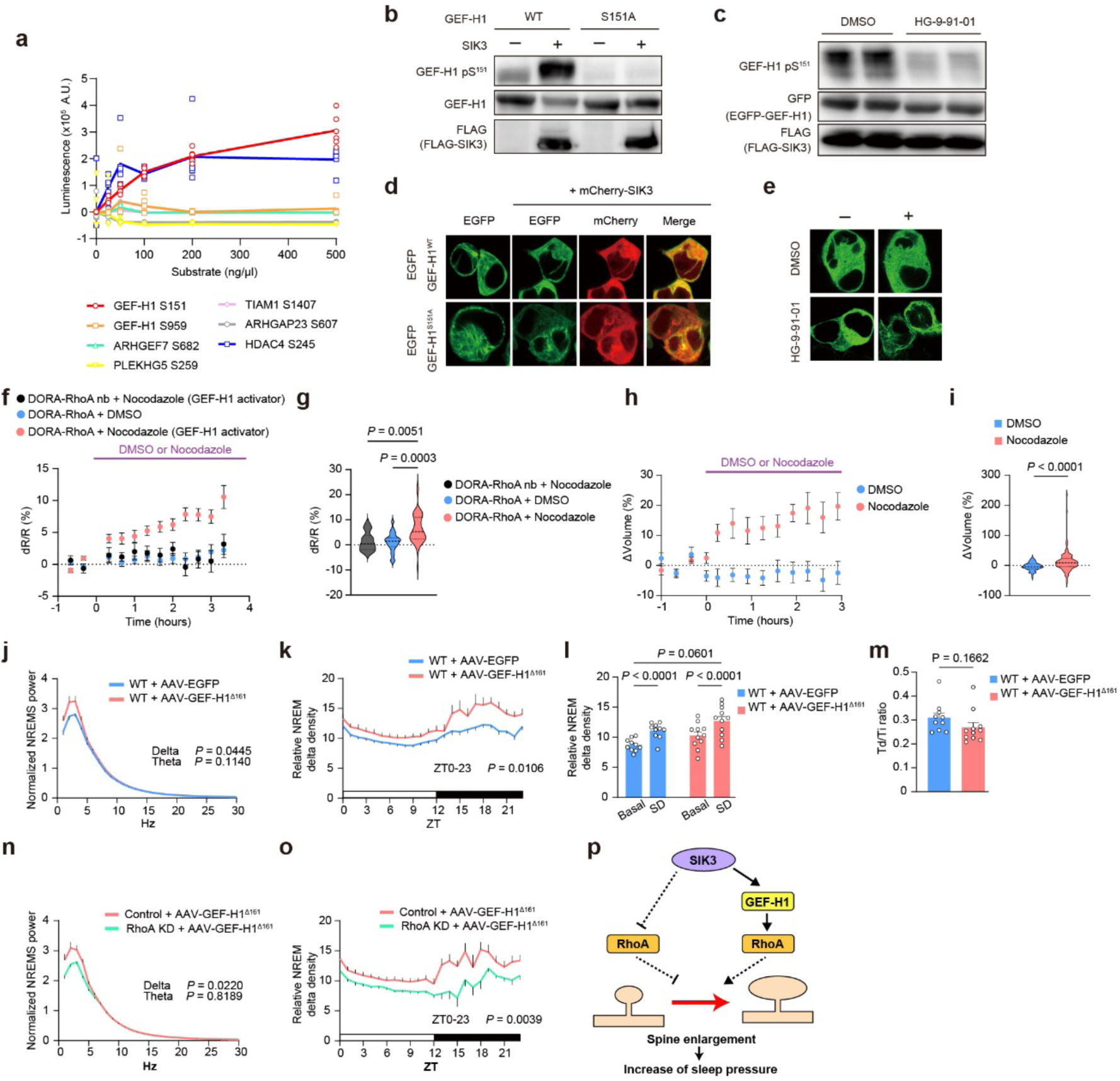
SIK3-GEF-H1-RhoA pathway enhances sleep pressure through spine enlargement. (A) In vitro kinase assay of SIK3 using GEF-H1 S151, S959, ARHGEF7 S682, ARHGAP23 S607, PLEKHG5 S259, TIAM1 S1407, and HDAC4 S245 peptide, which is a known SIK3 phosphorylation site. (B and C) Representative result of western immunostaining of HEK293T cells homogenates expressing GEF-H1 (B) and SIK3, and those treated with HG-9-91-01 (C). (D) Representative live images of HEK293T cells expressing EGFP-GEF-H1^WT^ or EGFP-GEF-H1^S151A^ alone or in combination with mCherry-SIK3. (E) Representative live images of HEK293T cells expressing EGFP-GEF-H1^WT^, treated with DMSO or HG-9-91-01. (F) Temporal FRET changes in RhoA activity during nocodazole stimulation. (G) The average dR/R from 0 to 200 min after Nocodazole administration (DORA-RhoA nb + Nocodazole; n (dendrite, neuron) = 14, 9, DORA-RhoA + DMSO; n (dendrite, neuron) = 23, 9, DORA-RhoA + Nocodazole; n (dendrite, neuron) = 51, 14). (H) The time course of changes in spine volume in the presence or absence of nocodazole. Delta are plotted as a percentage change from the baseline (1 hour before drug administration) average (DMSO; n (spine, neuron) = 67, 7, Nocodazole; n (spine, neuron) = 106, 5, U = 2204). (I) The average delta after 20 min after drug administration. (J and K) EEG power spectra during NREMS (J) and time course of NREMS delta density over 24 hours (K) (EGFP, n = 13; GEF-H1Δ161, n = 13). (L) NREMS delta density during ZT6–ZT8 under the basal condition and after 6-hour sleep deprivation (EGFP, n = 8; GEF-H1^Δ161^, n = 11). (M) The Td/Ti ratio derived from Process S fitting for sleep pressure dynamics (EGFP, n = 8; GEF-H1^Δ161^, n = 11). (N and O) EEG power spectra during NREMS (N), and time course of NREMS delta density over 24 hours (O) of WT mice given AAV.B10-GEF-H1^Δ161^ and AAV.B10-RhoA shRNA or AAV.B10-RhoA scramble shRNA (scramble shRNA, n = 8; RhoA shRNA, n = 11). (P) The proposed role of SIK3-RhoA pathway in the regulation of sleep pressure. SIK3 suppresses RhoA, thereby preventing RhoA from inhibiting spine enlargement, which consequently leads to an increase in spine size and an increase in sleep pressure. On the other hand, there is also a pathway in which SIK3 increases spine size and sleep pressure by promoting RhoA-mediated spine enlargement through the activation of GEF-H1. Dashed lines represent unknown mechanisms. Statistical significance was determined by Mann-Whitney U test (I), one-way ANOVA with Tukey’s test (G), unpaired t-test (J, M, N), two-way ANOVA with mixed-effects model (K, O), or two-way ANOVA with Tukey’s test (L). All error bars correspond to mean ± SEM.

### GEF-H1, a SIK3 substrate, positively regulates sleep pressure through RhoA

SIK3-KIOSS identified five proteins involved in the Rho GTPase signaling pathway as candidate SIK3 substrates (Figure 2B), suggesting that SIK3 may modulate RhoA activity through phosphorylation of upstream regulatory proteins. To determine which of these candidates are directly phosphorylated by SIK3, we examined the candidate sites, including GEF-H1 S151, S959, ARHGEF7 S682, PLEKHG5 S259, TIAM1 S1407, and ARHGAP23 S607, using a cell-free kinase assay with recombinant SIK3 and short candidate-site peptides. The GEF-H1 S151 peptide was phosphorylated by SIK3, whereas the other candidate-site peptides showed no detectable phosphorylation (Figure 4A). Consistent with this, co-expression of GEF-H1 with SIK3 in HEK293T cells markedly increased phosphorylation at S151 (Figure 4B). This signal was abolished by alanine substitution at S151 (GEF-H1^S151A^) and was significantly reduced by HG-9-91-01 treatment (Figure 4B and 4C). GEF-H1 is normally sequestered in an inhibited state on microtubules and phosphorylation at S151 promotes its dissociation from microtubules and catalytic activation^32,33^. When co-expressed with SIK3, a filamentous distribution of GEF-H1 consistent with microtubule association was disrupted, whereas the S151A substitution and HG-9-91-01 treatment inhibited this diffusion even in the presence of SIK3 (Figure 4D and 4E). Together, these results identify GEF-H1 as a direct substrate of SIK3 and suggest that SIK3 activates GEF-H1 through S151 phosphorylation.

Given that GEF-H1 is a direct SIK3 substrate, we asked whether GEF-H1 influences spine morphology and NREMS delta density. Here, we were faced with an obvious paradox: on the one hand, because GEF-H1 is well known as a Rho GEF that activates RhoA^34,35^, we predicted that SIK3 should activate RhoA through the activation of GEF-H1; on the other hand, our results described above (Figure 1-3) indicated that SIK3 enlarges spines and enhances NREMS delta density by inhibiting RhoA, which then acts as a negative regulator of spine size and NREMS delta density (Fig.4p). We administered nocodazole^36^, which can release GEF-H1 from microtubules and activate GEF-H1– RhoA signaling, to cultured cortical neurons in order to verify the function of GEF-H1. Whereas we confirmed that RhoA activity was enhanced upon nocodazole treatment (Figure 4F and 4G), we simultaneously found that nocodazole treatment increased spine size (Figure 4H and 4I). To investigate the effect of GEF-H1 on sleep pressure, we virally expressed a constitutively active form of GEF-H1 (GEF-H1^Δ161^) pan-neuronally in wild-type mice^37,38^. Consistent with the spine enlargement induced by GEF-H1 activation in vitro, EEG delta density during NREMS was increased in GEF-H1^Δ161^–expressing mice (Figure 4J and 4K), while GEF-H1^Δ161^ expression had no effect on wake and sleep amount or NREMS episode duration (Figure S5A-S5D). EEG delta density during REMS was also significantly increased in GEF-H1^Δ161^ expressing mice (Figure S5F). Neither the increase in NREMS delta density after 6 hours of sleep deprivation nor the inferred dynamics of Process S differed from controls (Figure 4L and 4M, Figure S5G and S5H). Next, to determine whether the effect of GEF-H1^Δ161^ on NREMS delta density requires RhoA, we neuronally co-expressed RhoA shRNAs with GEF-H1^Δ161^ throughout the brain. Although RhoA knockdown alone had no detectable effect on sleep phenotype (see above; Figure S3), it completely suppressed the increase in NREMS delta density induced by GEF-H1^Δ161^ (Figure 4N and 4O, Figure S6). These results indicate that the increase in NREMS delta density induced by GEF-H1^Δ161^ requires RhoA. Thus, in contrast to the SIK3-dependent reduction of RhoA activity observed during cLTP in cultured neurons, GEF-H1 activation promotes spine enlargement and increases NREMS delta density through a RhoA-dependent mechanism (Figure 4P).

## Discussion

Here, we showed that SIK3 links synaptic regulation to NREMS delta density through RhoA-mediated signaling based on three principal observations: SIK3 promotes activity-dependent spine enlargement; RhoA signaling modulates both spine size and NREMS delta density; and GEF-H1 is a direct SIK3 substrate whose effect on NREMS delta density requires RhoA. Notably, however, the direction of RhoA-dependent effects differed across experimental contexts. During cLTP in cultured neurons, RhoA activity in spines was reduced as spines enlarged. In contrast, activation of GEF-H1 promoted both spine enlargement and increased NREMS delta density through RhoA-dependent mechanisms. Thus, the relationship among RhoA activity, spine morphology and NREMS delta density is context dependent rather than unidirectional.

Such context-dependence is consistent with the known spatial and temporal organisation of Rho-family GTPase signalling. Although RhoA has often been associated with spine destabilisation and shrinkage^39^, RhoA can also support spine enlargement under specific conditions^40–42^. The outcome of RhoA signalling is shaped by upstream regulatory molecules, subcellular localisation, and interactions with other Rho-family GTPases, including Rac1 and Cdc42^43–45^. In the present study, the cLTP procedure recruits synaptic NMDAR-dependent Ca²⁺ signalling and actin-regulatory pathways such as the Rac/PAK/LIMK cascade^22,46^, whereas nocodazole activates GEF-H1–RhoA signalling through disruption of microtubules^36^. These distinct signalling contexts likely account for the different structural outcomes associated with RhoA activation^47^.

Our results suggest that SIK3 can influence RhoA signalling in opposing directions depending on the upstream stimulus and downstream effector context (Fig.4p). SIK3-dependent phosphorylation of GEF-H1 should promote RhoA activation, whereas we observed that the cLTP procedure reduces RhoA activity in an SIK3-dependent manner. Thus, SIK3 may engage a distinct combination of Rho GEFs and Rho GAPs to shape the spatial and temporal profile of RhoA signalling. The SIK3-KIOSS results suggested that both Rho GEFs and Rho GAPs are (direct and indirect) targets of SIK3, and many Rho GEFs and Rho GAPs possess multiple signal transduction domains within their molecules and can recruit multiple downstream factors^48,49^. Based on these findings, it is possible that SIK3 selectively utilises Rho GEFs and Rho GAPs to modulate the appropriate downstream pathways for each context, thereby eventually promoting spine enlargement.

The current study also suggests that NREMS amount and NREMS delta density are independently regulated. Sleep pressure increases with extended wakefulness and is attenuated by sleep, indicating that sleep pressure is regulated by the history of sleep/wake^9^. However, multiple studies have shown that NREM delta density does not simply correlate with total sleep amount^2,50,51^. Furthermore, previous studies indicate that the augmentation of NREMS delta density concomitant with escalating sleep pressure occurs predominantly within the prefrontal cortex^2,50,52^, and SIK3 modulates NREMS delta density primarily in the cerebral cortex^11^. In this study, constitutive activation of RhoA and GEF-H1 altered NREMS delta density without a significant effect on NREMS amount, supporting the view that sleep amount and delta density are independent factors. Our findings place SIK3 at the interface between kinase-dependent sleep regulation and the cytoskeletal mechanisms underlying synaptic plasticity. Rather than acting through a single linear pathway, SIK3 may recruit context-dependent Rho GTPase regulatory modules to adjust dendritic spine morphology and delta density during NREMS. This framework provides a complex molecular basis for coupling synaptic plasticity to the homeostatic regulation of sleep.

Previous studies have indicated that Liver kinase B1 (LKB1), cyclic AMP-dependent protein kinase A (PKA), and calcineurin function as upstream regulators of SIK3 in the sleep regulation pathway^11,12,53–55^. The timing of SIK3-RhoA pathway activation is presumably determined by these upstream signals. In addition, it has been shown that transcriptional regulators including HDAC4, HDAC5, and CREB-regulated transcription coactivator 1 (CRTC1), regulate NREMS delta density downstream of SIK3^11,12,56^. Based on these findings, it is likely that multiple independent mechanisms controlling sleep pressure, including synaptic regulation and gene expression control, are at work downstream of SIK3. Understanding how sleep pressure emerges from cellular-level processes, including the SIK3 pathways, will be key to revealing the fundamental function of sleep, providing insight into sleep disorder mechanisms, and shaping the next generation of sleep therapeutics.

## Resource availability

The mass spectrometry datasets, including raw data files and search engine files, have been deposited to MassIVE with accession code MSV000102488.

## Acknowledgements

We thank all members of the Yanagisawa/ Funato Lab, Shi Lab and the International Institute for Integrative Sleep Medicine, especially K. Ebihara and T. Kanae for technical support and K. Sakurai and Y.Ishikawa for technical comments. This work was supported by the World Premier International Research Center Initiative from MEXT to M.Y., AMED (JP21zf0127005 to M.Y.); JSPS KAKENHI (17H06095, 22H04918, 22K21351 to M.Y., H.F; 20H00567, 22K19410, 24H00671, 24H01219 to H.F.; 26507003, 18968064 to C.M., H.F.; 19K21206, 21K15339, 24K18213 to T.K.), JSPS PD (22KJ0337 to T.K), JSPS DC2 (25KJ0691 to M.J.), JST SPRING (HIJ07009 to M.J.) and JST CREST (JPMJCR1655 to M.Y.), JST Grant Number JPMJPF2210 and JPMJSP2124. Funding Program for World-Leading Innovative R&D on Science and Technology (FIRST Program) from JSPS to M.Y. Research Grants for the Medical Frontier Conference (TMFC) to T.K.

## Author contributions

T.K., M.J., and K.N. contributed equally to this work. T.K. performed KIOSS and the following kinase assay, and all sleep analyses. M.J., K.N., S. Nomura, and C.S. conducted the in vitro cLTP experiments under the guidance of T.S. and S.S. M.J. conducted the in vitro pharmacological experiments under the guidance of T.S. and S.S. M.J. and C.S. performed in vivo spine imaging, including virus preparation, virus injection, imaging procedures, and the data analysis, with S. Nomura, Y.I., T.S., and S.S. providing guidance. M.J. introduced cultured cortical neuron experiments with N.A-I. providing guidance. K.N. and A.K. performed EGFP-GEF-H1 imaging analysis. A.K. and N.H. performed western blotting. K.M. and S. Nakata developed the AAV.B10 retro-orbital injection system. T.K., M.J. and K.N. wrote the manuscript. M.Y., H.F. and S.S. conceived and supervised the entire project.

## Methods

### Animals

Mice were housed under controlled temperature and humidity conditions and maintained on a 12-h light: 12-h dark cycle. All animal experimental procedures were approved (Approved protocol ID 21-313) and conducted following the guidelines established by the Institutional Animal Care and Use Committee of the University of Tsukuba. Wild-type mice (C57BL/6N) were obtained from CLEA Japan and female ICR mice were obtained from Jackson Laboratory.

### Sleep analysis

EEG/EMG electrode implantation, EEG/EMG signal acquisition, and data analysis were performed as previously described^57,58^. At 7-9 weeks of age, male mice were implanted with EEG/EMG electrodes under anaesthesia with isoflurane. One week after the implantation, AAV.B10 was administered by retro-orbital injection under 5 % isoflurane anaesthesia. One week after the retro-orbital injection, mice were attached to a tether cable with a counterbalanced arm for habituation to the recording conditions for at least 1 week before EEG–EMG recording.

Basal EEG and EMG recordings were taken for two consecutive days from ZT0. Mice were sleep-deprived for 6 h from light-on (ZT0) immediately following the basal recording by gentle handling. EEG signals were subjected to fast Fourier transform analysis from 1 to 30 Hz with a 1-Hz bin using MATLAB. For the power spectrum of sleep–wakefulness, the EEG power of each frequency bin was expressed as a percentage of the mean total EEG power over the sum of 16–30 Hz and sleep–wake states.

### AAV production and administration

Adeno associated viruses (AAVs) were generated following previously reported procedures^11^. Briefly, AAVpro 293T cells (Takara Bio #632273) were co-transfected with plasmids encoding the AAV vector, pHelper, and pUCmini-iCAP-AAV.CAP-B10 (Addgene #175004). Seventy-two hours after transfection, cells were collected, and AAV particles were purified by iodixanol gradient ultracentrifugation. Viral genome titres were determined by qPCR with Thunderbird SYBR qPCR mix (TOYOBO #QPS-201). The AAV vector used to overexpress GEF-H1^Δ161^ in our study, pAAV[Exp]-SYN1>mArhgef2[NM_008487.4]*: WPRE, was constructed and packaged by VectorBuilder Inc. (Chicago, IL, USA). The vector ID is VB250416-1893nqh, which can be used to retrieve detailed information about the vector on vectorbuilder.com. The shRNA sequences were designed based on The RNAi Consortium (TRC). The RhoA shRNA sequences used are from TRC clone IDs TRCN0000304690, TRCN0000302388, and TRCN0000304743, and the scrambled shRNA sequences were obtained by rearranging these sequences.

### Cell Culture (HEK293T cells)

HEK293T cells (RCB2022, the RIKEN BRC Cell Bank) were cultured in Dulbecco’s modified Eagle’s medium (DMEM) with 10% FBS and penicillin/streptomycin. For plasmid transfection, HEK293T cells were grown to 80% confluency in 10 cm plates.

### Western blotting

Western blotting was performed as previously described^10,59^. The following antibodies were used: anti-HDAC4 (D15C3, CST #7628), anti-Phospho-HDAC4(Ser246)/HDAC5(Ser259)/HDAC7(Ser155) (D27B5, CST #3443), anti-phospho-PKA substrate (RRXS/T) (CST #9624), anti-β-tubulin (9F3, CST #2128), anti-FLAG (Wako 012-22384), anti-GFP (Santa Cruz sc-9996), and anti-GEF-H1 (COSMO BIO, 24472-1-AP). A rabbit polyclonal antibody against the phosphorylated Ser151 of mouse GEF-H1 was generated using a custom antibody production service (Eurofins). One day before cell lysate extraction, HEK293T cells were transfected with 16 μg of plasmid DNA using 70 μL of polyethylenimine MAX (Cosmo Bio #24765-1). HEK-293T cells were homogenised in cold Buffer C (20 mM HEPES-Na, pH.7.5, 100 mM NaCl, 1.5 % Triton-X100, 15 mM NaF, 10 mM Na_4_P_2_O_7_, and 5 mM EDTA) supplemented with protease inhibitor (Roche) and phosphatase inhibitor (Roche). The cell lysates were rotated for 20 min at 4 °C and centrifuged at 14,000 g for 12 min at 4 °C. The supernatant was used for western blotting.

### Kinase activity assay

In vitro kinase activity of SIK3 was measured using an ADP-Glo Kinase Assay kit (Promega) with recombinant active SIK3 (SignalChem #S12-11G). The peptide substrates for active SIK3 were GEF-H1tide (Ser151: SLAKSVSTTNIAGRRR, mouse GEF-H1 145-157 with an additional three arginine residues at the C-terminal end; Ser959: RLSPPPHSPRDFTRRRR, mouse GEF-H1 953-965 with additional three arginine residues at the C-terminal end), ARHGEF7tide (Ser682: EFAVRKSTAALEERRR, mouse ARHGEF7 676-688 with an additional three arginine residues at the C-terminal end), TIAM1tide (Ser1407: QLLKTESLPSAQQRRR, mouse TIAM1 1401-1413 with an additional three arginine residues at the C-terminal end), PLEKHG5tide (Ser259: PQSRRESSLDILARRR, mouse PLEKHG5 253-265 with an additional three arginine residues at the C-terminal end), ARHGAP21tide (Ser469: MKYCPRSASQGALRRR, mouse ARHGAP21 463-475 with an additional three arginine residues at the C-terminal end), and ARHGAP23tide (Ser607: KAGRRSSYLLAITRRR, mouse ARHGAP23 601-613 with an additional three arginine residues at the C-terminal end).

### Mass spectrometry

Based on the Kinase-Oriented Substrate Screening (KIOSS) method described by Nishioka et al^60^. Confluent N41 cells in 10-cm dishes were cultured with FBS free D-MEM (Wako) overnight in a 37 °C, 5 % incubator. The next day, 10 μM HG-9-91-01 or 5 % DMSO in FBS free D-MEM was added to the cells, and the cells were incubated for 30 min in a 37 °C, 5 % incubator. Then, 250 nM calyculin A or 2.5 % DMSO in FBS free D-MEM was added to the cells, and the cells were incubated for 10 min in a 37 °C, 5 % incubator. Cells were lysed in 1 mL lysis buffer [20 mM Tris/HCl, 150 mM NaCl, 1 mM EDTA, 1 mM DTT, 1 % Igepal CA-630 (Sigma), protease inhibitor cocktail (Roche, Indianapolis, IN), and PhosStop (Roche), pH 7.5] through 15 s of sonication. The lysate was centrifuged at 20,000×g for 10 min at 4°C; the supernatant was used as the cell lysate. A total of 500 pmol of GST-14-3-3ζ, immobilised on glutathione Sepharose 4B beads, was incubated with the cell lysate for 1 h at 4°C with rotation. The beads were then washed three times with wash buffer (20 mM Tris/HCl, 150 mM NaCl, 1 mM EDTA, 1 mM DTT, and 1 % NP-40, pH 6.8) and an additional three times with detergent-free buffer (20 mM Tris/HCl, 150 mM NaCl, 1 mM EDTA, and 1 mM DTT, pH 6.8). The bound proteins were extracted from the beads using guanidine solution (7 M guanidine and 50 mM Tris, pH 8.0), reduced through incubation in 5 mM dithiothreitol for 30 min, and alkylated using 10 mM iodoacetamide for 1 h in the dark. The proteins were demineralised, concentrated through methanol/chloroform precipitation and digested with trypsin (50 mM NH4HCO3, 1.2 M urea, and 0.5 µg trypsin). Phosphopeptide enrichment was performed using a Titansphere Phos-TiO Kit (GL Sciences, Tokyo, Japan), followed by demineralisation using SPE c-tips (Nikkyo Technos, Tokyo, Japan) according to the manufacturer’s instructions. LC/MS/MS was performed using the Orbitrap-Fusion mass spectrometry platform coupled with EASY-nLC 1000 liquid chromatography pump (Thermo Fisher Scientific). A pre-column (Acclaim PepMap 100 C18, Thermo Fisher Scientific) and analytical column (NTCC-360/75-3-125, NIKKYO) were used for sample trapping and analytical separation. The peptides were separated using a 70-min gradient of 5 %–40 % acetonitrile containing 0.1% formic acid at a flow rate of 400 nL/min. A peak list was generated and calibrated using Proteome Discoverer software (PD, v.2.1, Thermo Fisher Scientific). The target mouse protein database was generated from UniProt (17 October 2015). MS^2^ spectra were searched with ± 20 ppm for precursor ion mass tolerance, ± 1 Da for fragment ion mass tolerance, fully tryptic restriction, four maximal missed cleavages, dynamic mass shift for oxidation of methionine (+15.9949 Da), fixed TMT modifications on the N terminus and lysine (+229.1629 Da), and carbamidomethylation of cysteine residues (+57.0215 Da). For phosphoproteomic analysis, additional dynamic modifications on serine, threonine and tyrosine (+79.9663 Da) were used. The peptide spectrum matches (PSMs) were filtered by Percolator32 (PD 2.1) to achieve 1% protein and peptide FDR (according to Q value) for proteome and phosphoproteome, respectively. ptmRS33 (PD 2.1) was used for phosphorylation site localisation, which derived a localisation probability score for each putatively modified site based on the given MS2 data. Phosphopeptides with phosphorylation site probability scores ≥ 25 were considered in the following analysis. Candidate sites were defined as those showing a fivefold or greater difference in intensity between calyculin A treatment and combined treatment with HG-9-91-01.

### Live imaging in HEK293T cells

3 days before observation, HEK293T cells at a density of 3.5×10^5^ cells/ml were plated in 35-mm glass bottom dishes coated with 0.04% Polyethyleneimine. The following day, cells were co-transfected with the plasmids encoding EGFP-GEF-H1 or mCherry-SIK3 using Lipofectamine LTX reagent (Thermo Fisher Scientific, Waltham, MA, #15338100) and incubated for 2 days at 37℃, 5% CO_2_. Before observation, DMEM with serum was replaced with phenol red-free FluoroBrite DMEM (Thermo Fisher Scientific, Waltham, MA, A1896701) warmed to 37°C. Cells were observed using the confocal microscope Observer Z1 (Carl Zeiss, Oberkochen, Germany), equipped with an objective lens (Objective Plan-Apochromat 63x/1.4 Oil M27; Carl Zeiss), an FC12 definite focus autofocusing system (Carl Zeiss), and an INUG2TF stage-top incubator (Tokai Hit, Fujinomiya, Japan). Cells were incubated during observation at 37 °C, 5% CO_2_. For pharmacological experiments, cells were treated with HG-9-91-01 or DMSO after baseline observation.

### Analysis of in vivo spine size

To achieve sparse labeling of cortical neurons with mScarlet, a double-floxed inverted open reading frame (DIO) strategy was combined with low-level Cre expression. For viral delivery, AAV vectors encoding CaMKII(0.4)-Cre (1.0 × 10⁹ genome copies (GC)/ml) and hSyn-DIO-mScarlet (1.0 × 10¹³ GC/ml) were mixed, and approximately 0.5 µL of a viral solution was injected at a rate of 0.1 µL /min into the PFC (AP +1.5 mm, ML ±0.6 mm, DV +2.0 mm) from bregma. Animals were deeply anesthetized with isoflurane (Wako #099-06571), transcardially perfused with 4% paraformaldehyde (Nacalai Tesque #26126-25) at ZT12, and decapitated. Brains were extracted and coronally sectioned at a thickness of 40 µm using a cryostat CM3050 (Leica, Wetzlar, Germany). To enhance mScarlet signal detection, sections were immunostained with an anti-RFP primary antibody (1:1,000; MBL, PM005), followed by a goat anti-rabbit IgG Alexa Fluor 594–conjugated secondary antibody (1:400; Jackson ImmunoResearch, #111-587-008). Dendritic spines were imaged using a confocal microscope TCS SP8 (Leica, Wetzlar, Germany). Z-stack images were acquired with a step size of 0.5 µm. Image segmentation of dendrites and spines was performed using DeepD3^61^ with pretrained models. Three-dimensional regions of interest (ROIs) corresponding to individual spines were generated based on the segmentation output. For each spine ROI, voxel values were calculated by summing pixel intensities along the z-axis. Voxel values were averaged for each dendrite and used for subsequent statistical analyses. Sample sizes were not predetermined prior to experimentation. To assess whether DeepD3 can be applied to Sleepy mice datasets, we compared spine annotations generated by manual labeling with those produced by DeepD3, and calculated precision and recall based on the number of detected spines. DeepD3 can recognize mutant mice with the same accuracy as wild-type mice (Figure S1).

### Imaging of spines in cultured cortical neurons

Cultured cortical neurons were prepared from embryonic day 17 (E17) ICR mouse embryos. Cortical tissues were dissociated and neurons were seeded onto poly-L-lysine–coated glass-bottom dishes (Iwaki #5816-006) at a density of 5.0 × 10⁵ cells/cm². Cells were maintained at 37 °C in a humidified incubator with 5% CO₂ using Neurobasal medium supplemented with B27 and L-glutamine (Nacalai Tesque #16948-04). Neurons were transfected with a CAG-driven mScarlet construct at 13–14 days in vitro (DIV) using Lipofectamine 2000 (Invitrogen #11668019), according to the manufacturer’s protocol. Time-lapse confocal imaging was performed at DIV19–21 using Observer Z1, equipped with an objective lens (Objective Plan-Apochromat 63x/1.4 Oil M27). Images were acquired at 20-minute intervals. Chemical long-term potentiation (cLTP) was induced as follows. Neurons were pretreated with 50 µM D-APV (CAY #14539) for 12 hours prior to stimulation. The culture medium was then replaced with a stimulation solution containing 140 mM NaCl (Nacalai Tesque #31320-05), 5 mM KCl (Nacalai Tesque #28514-75), 10 mM D-glucose (Nacalai Tesque #16806-25), 2 mM CaCl₂ (Nacalai Tesque #06729-55), 10 mM HEPES-Na, 200 µM glycine (Wako #077-00735), and 20 µM bicuculline (abcam #ab144491). After 15 minutes of incubation, the stimulation solution was replaced with the original culture medium. Nocodazole (MilliporeSigma #M1404) was added at 200 nM after baseline. Spine volume was calculated by the method previously described^2^.

### FRET imaging

Cultured neurons at 13–14 days in vitro (DIV) were transfected with apCAGGS-DORA-RhoA or the RhoA binding-deficient control construct pCAGGS-DORA-RhoA-nb, which contains the PKN1(L59Q) mutation^29,30^. Both constructs were kindly provided by Dr. Imai. For FRET imaging, neurons were excited with a 445 nm laser, and emission signals were collected through a short-pass filter (< 490 nm) for the donor channel and a band-pass filter (530–555 nm) for the FRET channel. The imaging time course and drug-treatment protocol were identical to those used for spine imaging. Z-sum projections of FRET images were corrected using the Registration plugin (Correct 3D drift) and subjected to background subtraction in ImageJ. ROIs were manually defined for individual spines, and CFP and YFP intensity were calculated in the spines. Relative changes in FRET ratio (*dR*/*R*) were calculated as follows:

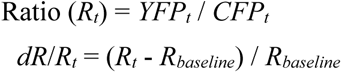

where (*R_t_*) is the YFP/CFP ratio at time point (t), and R_baseline_ is the mean YFP/CFP ratio during the baseline.

### Statistical analysis

The experimenters were blinded to genotypes. Statistical analyses were performed using Prism 10 (GraphPad). For two group comparison, unpaired t-test or Mann-Whitney U test was used for data with or without a normal distribution, respectively. We used two-way ANOVA for multiple comparisons involving two independent variables, and a mixed effect model for repeated measured data with missing values^62^. ANOVA analyses were subjected to the Tukey test. P < 0.05 was considered statistically significant. Data shown as mean ± standard error of mean (s.e.m). Two-sided P values were used.

---

**Figure S1.**
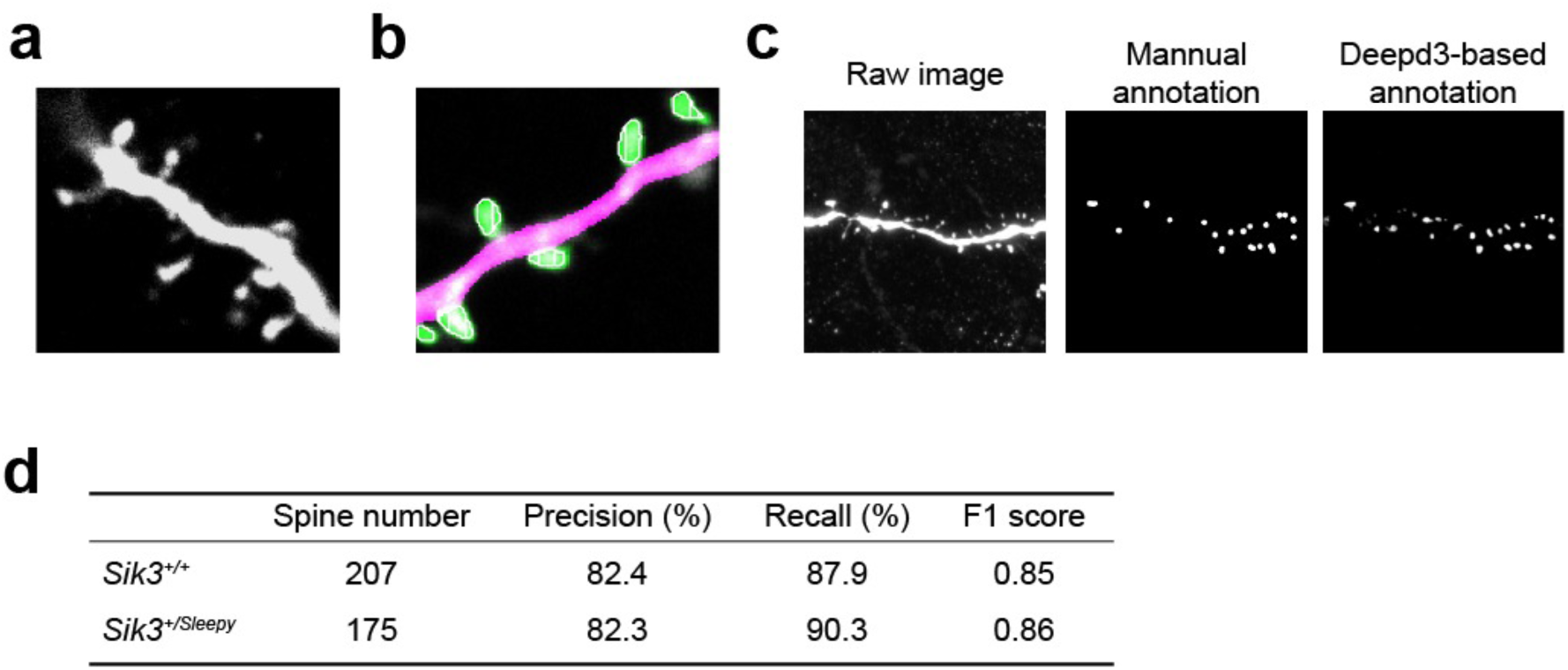
Representative images of spines in the PFC and performance metrics. (A) Representative images of spines in the PFC. Scale bars, 2.5 µm. (B) Representative images of automatic segmentation of the spines using DeepD3. Spines (yellow-green) and dendrites (pink) are shown. (C) Raw image (left), manual annotation of spines (center), and DeepD3-based annotation of spines (right). The raw image is a z-projection of the acquired z-stack. (D) Performance metrics (precision, recall, and F1 score) for model-based spine annotation relative to human annotations.

**Figure S2.**
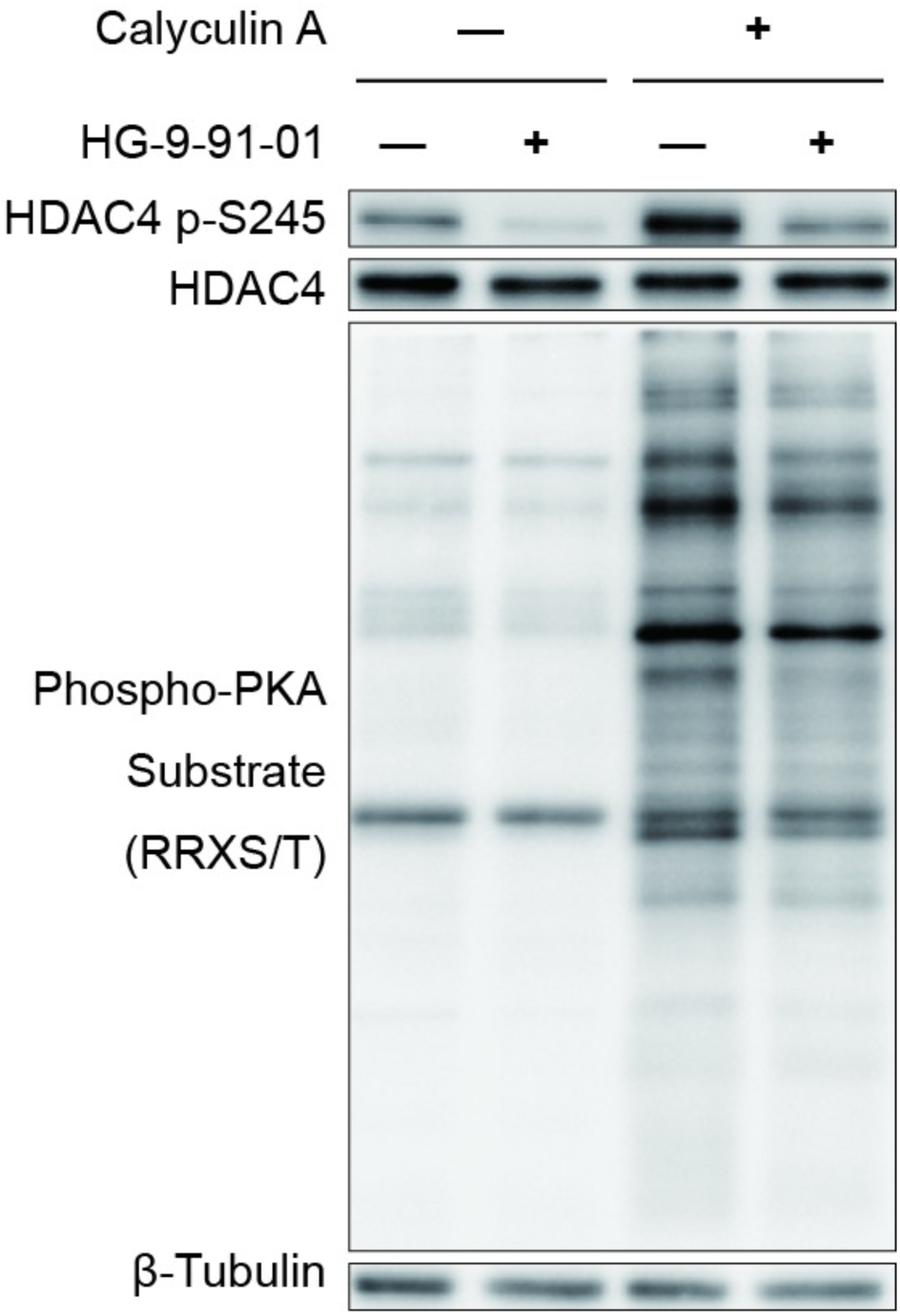
HG-9-91-01 specifically suppresses phosphorylation of SIK substrate. Representative result of western immunostaining of HEK293T cells homogenates treated with DMSO, HG-9-91-01, Calyculin A or HG-9-91-01 and Calyculin A.

**Figure S3.**
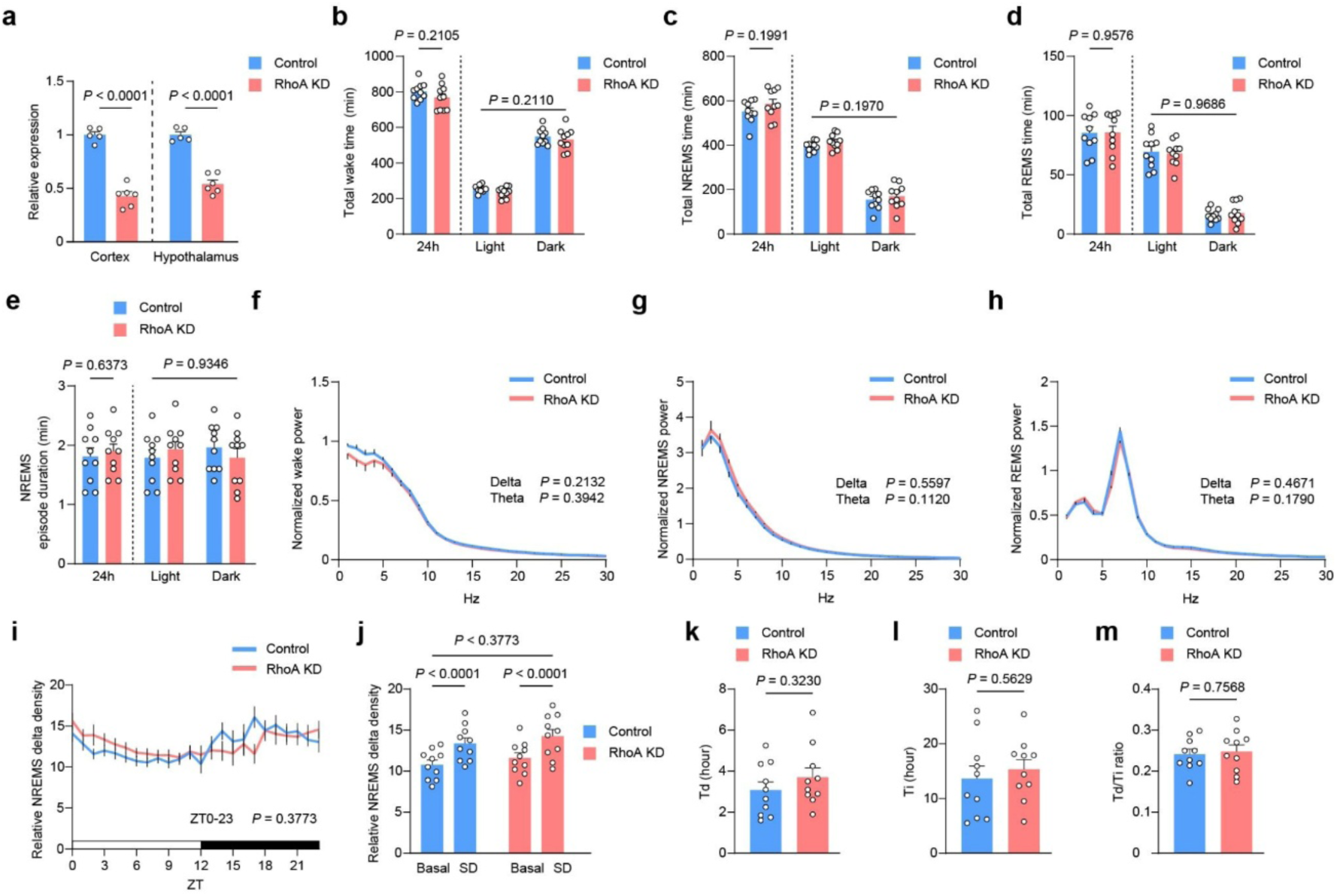
Sleep/wakefulness of RhoA knockdown mice. (A) RhoA mRNA expression in the cerebral cortex and hypothalamus after two weeks of retro-orbital injection (control, n = 5; Rhoa KD, n = 6). (B-E) Total time spent during wakefulness (B), NREMS (C), REMS (D), and NREMS episode duration (E) over 24 hours and during light and dark phases of EGFP expressing or Rhoa KD mice (control, n = 10; Rhoa KD, n = 10). (F-I) EEG power spectra during wakefulness (F), NREMS (G), REMS (H) over 24 hours and time course of NREMS delta density over 24 hours (I) (control, n = 10; Rhoa KD, n = 10). (J) NREMS delta density during ZT6–ZT8 under the basal condition and after 6-hour sleep deprivation (control, n = 10; Rhoa KD, n = 10). (K-M) *T*_d_ (K), *T*_i_ (L), and the *T*_d_/*T*_i_ ratio (M) derived from Process S fitting for sleep pressure dynamics (control, n = 10; Rhoa KD, n = 10). Statistical significance was determined by two-way ANOVA with Tukey’s test (B-E right, J), unpaired t-test (A, B-E left, F-H, K-M), or two-way ANOVA with mixed-effects model (I). All error bars correspond to mean ± SEM.

**Figure S4.**
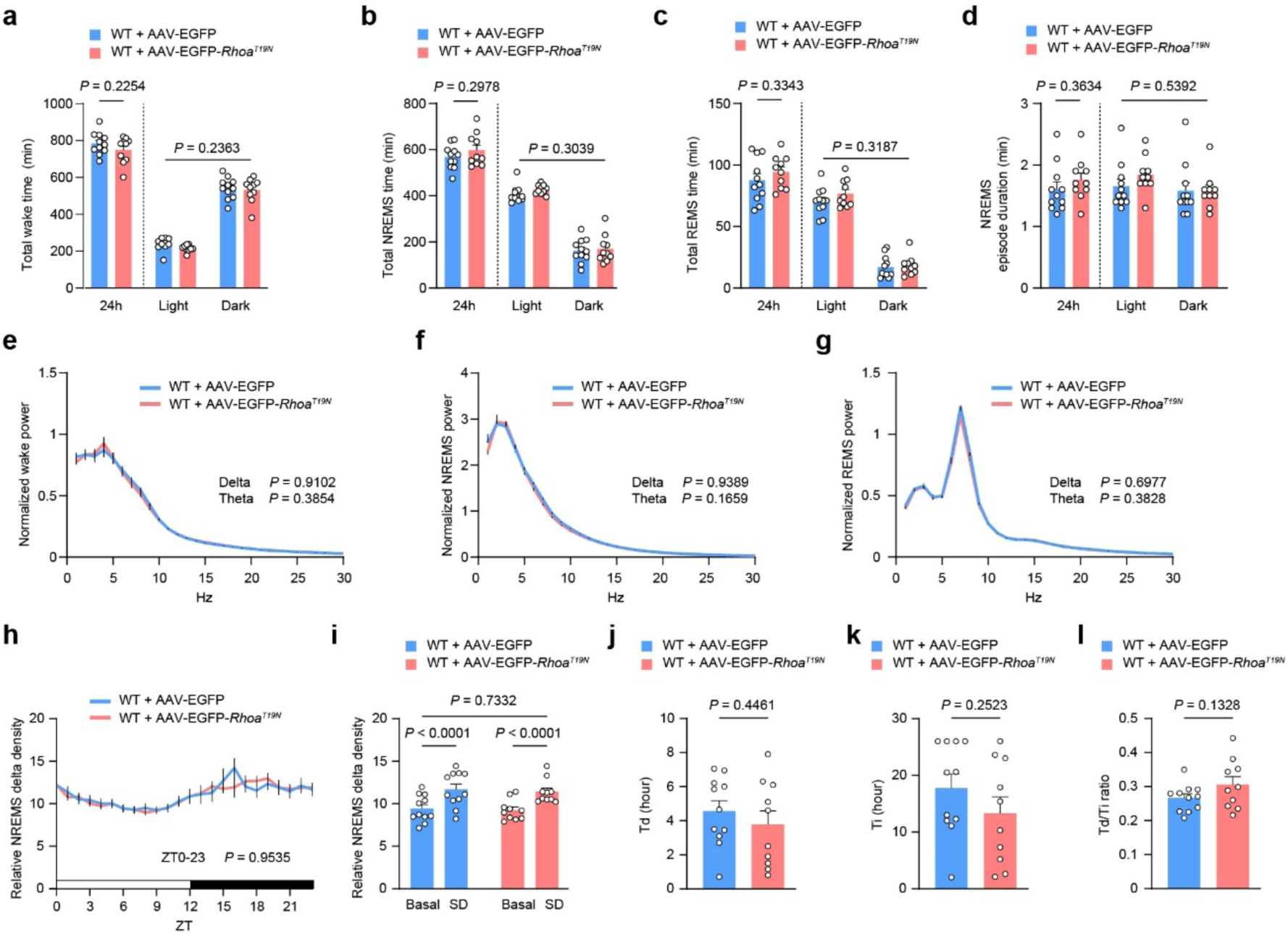
Sleep/wakefulness of dominant negative RhoA expressing mice. (A-D) Total time spent during wakefulness (A), NREMS (B), REMS (C), and NREMS episode duration (D) over 24 hours and during light and dark phases of WT mice given AAV.B10-EGFP or EGFP-*Rhoa^T19N^*(control, n = 11; *Rhoa^T19N^*, n = 10). (E-H) EEG power spectra during wakefulness (E), NREMS (F), REMS (G) over 24 hours (control, n = 11; *Rhoa^T19N^*, n = 10) and time course of NREMS delta density over 24 hours (H). (I) NREMS delta density during ZT6–ZT8 under the basal condition and after 6-hour sleep deprivation (control, n = 11; *Rhoa^T19N^*, n = 10). (J-L) *T*_d_ (J), *T*_i_ (K), and the *T*_d_/*T*_i_ ratio (L) derived from Process S fitting for sleep pressure dynamics (control, n = 11; *Rhoa^T19N^*, n = 10). Statistical significance was determined by two-way ANOVA with Tukey’s test (A-D right, I), unpaired t-test (A-D left, E-G, J-L), or two-way ANOVA with mixed-effects model (H). All error bars correspond to mean ± SEM.

**Figure S5.**
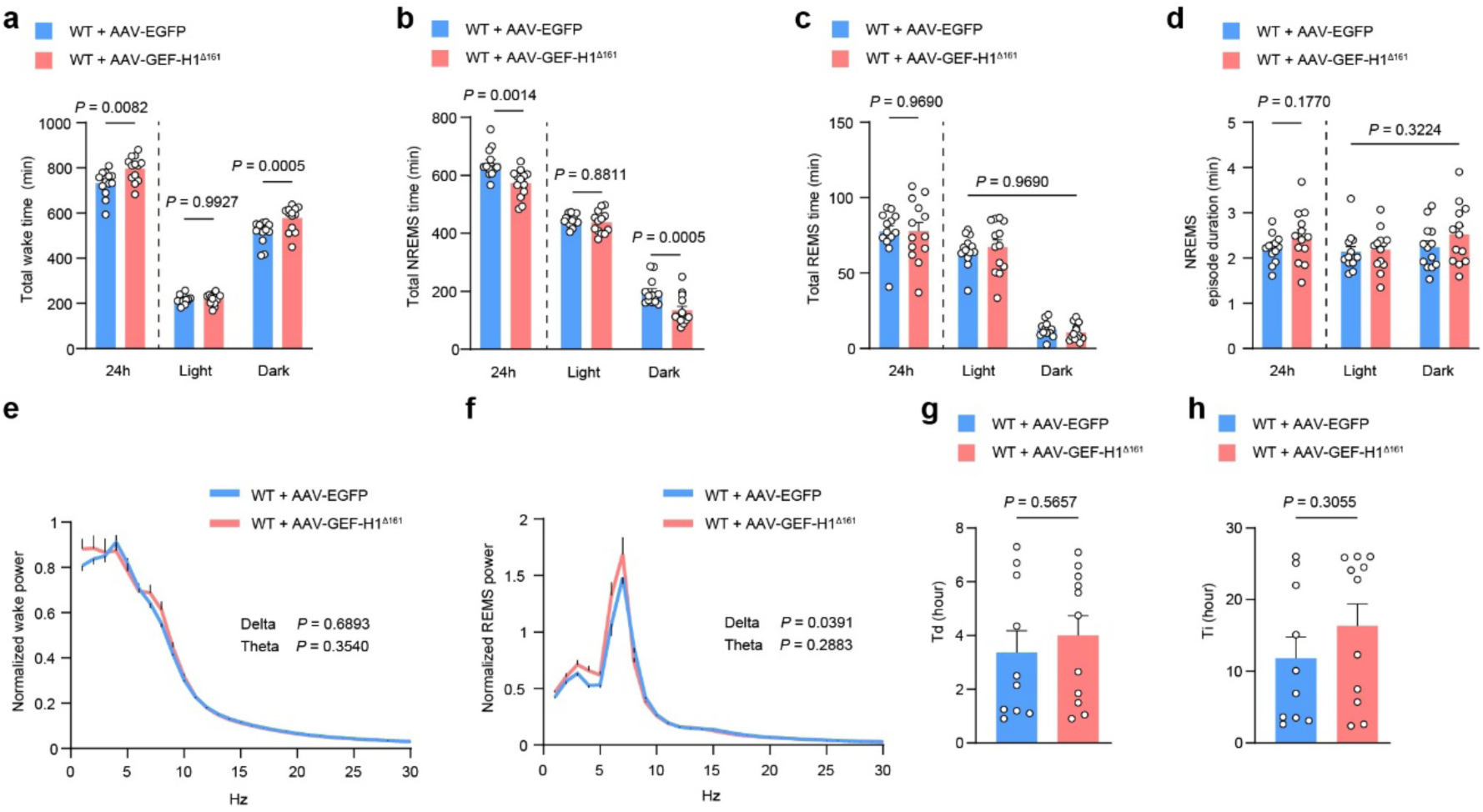
Sleep/wakefulness of constitutively active GEF-H1 expressing mice. (A-D) Total time spent during wakefulness (A), NREMS (B), REMS (C), and NREMS episode duration (D) over 24 hours and during light and dark phases of WT mice given AAV.B10-EGFP or GEF-H1^Δ161^ (EGFP, n = 13; GEF-H1^Δ161^, n = 13). (E and F) EEG power spectra during wakefulness (E) and REMS (F) (EGFP, n = 13; GEF-H1^Δ161^, n = 13). (G and H) Td (G) and Ti (H) derived from Process S fitting for sleep pressure dynamics (EGFP, n = 8; GEF-H1^Δ161^, n = 11). Statistical significance was determined by unpaired t-test (A-D left, E-H) or two-way ANOVA with Tukey’s test (A-D right). All error bars correspond to mean ± SEM.

**Figure S6.**
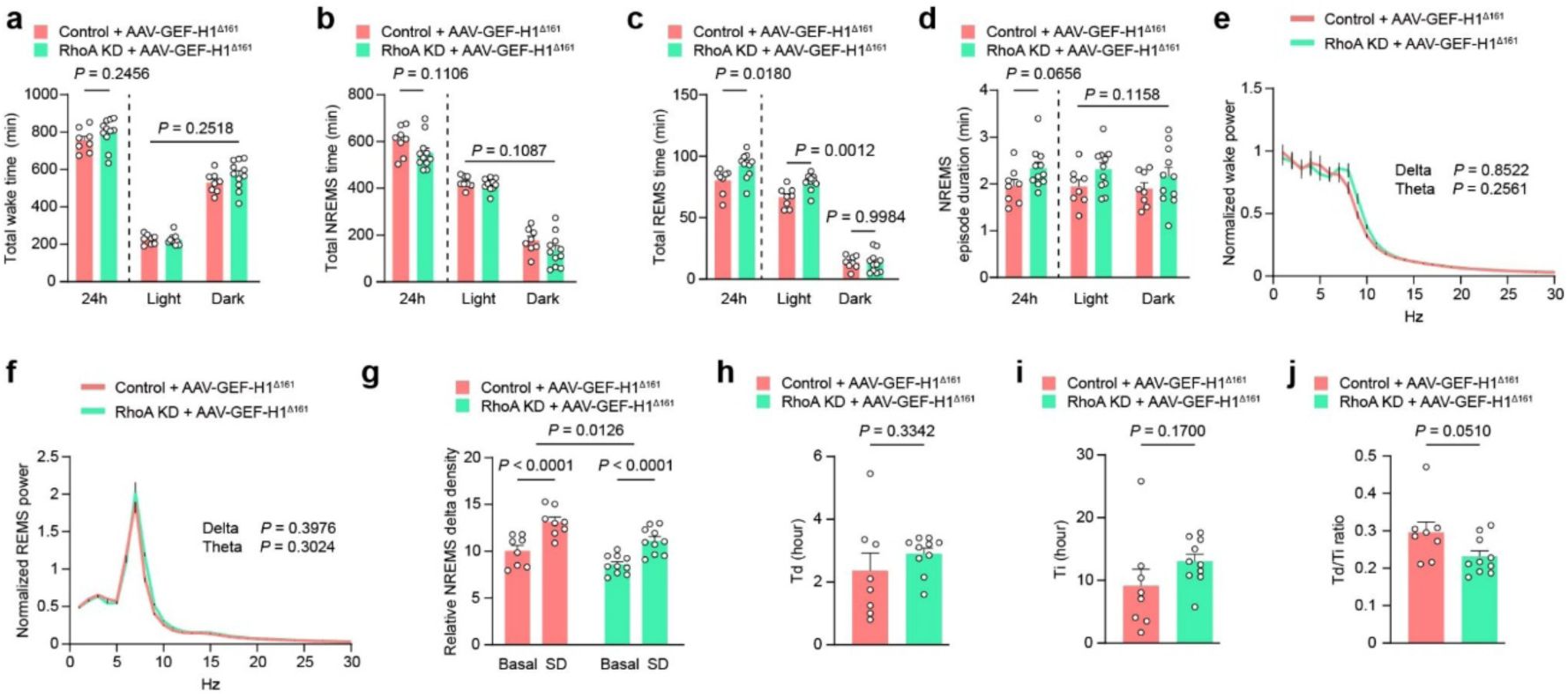
Sleep/wakefulness of constitutively active GEF-H1 expressing and Rhoa gene-modified mice. (A-D) Total time spent during wakefulness (A), NREMS (B), REMS (C), and NREMS episode duration (D) over 24 hours and during light and dark phases of control mice or Rhoa knockdown mice given AAV.B10-GEF-H1^Δ161^ (control, n = 8; Rhoa KD, n = 11). (E and F) EEG power spectra during wakefulness (E), REMS (F) over 24 hours (control, n = 8; Rhoa KD, n = 11). (G) NREMS delta density during ZT6–ZT8 under the basal condition and after 6-hour sleep deprivation (EGFP + GEF-H1^Δ161^, n = 8; RhoA KD + GEF-H1^Δ161^, n = 10). (H-J) Td (H), Ti (I), and the Td/Ti ratio (J) derived from Process S fitting for sleep pressure dynamics (EGFP + GEF-H1^Δ161^, n = 8; RhoA KD + GEF-H1^Δ161^, n = 10). Statistical significance was determined by unpaired t-test (A-D left, E, F, H-J) or two-way ANOVA with Tukey’s test (A-D right, G). All error bars correspond to mean ± SEM.

